# MotorBench: A Cryo-Electron Tomography Dataset of Bacterial Flagellar Motors for Testing Detection Algorithms

**DOI:** 10.1101/2025.04.23.650258

**Authors:** C. Braxton Owens, Rachel Webb, T. J. Hart, Matthew M. Ward, Andrew J. Darley, Stefano Maggi, Bryan S. Morse, Grant J. Jensen, Walter C. Reade, Mohammed Kaplan, Gus L.W. Hart

## Abstract

Understanding bacterial nanomachines like flagellar motors, which are crucial for pathogenic bacteria motility, is vital for microbiological and therapeutic research. Cryogenic electron tomography (cryo-ET) enables visualization of these structures within cells at near-native conditions. But manual identification remains challenging due to low contrast, limited resolution, and crowded *in vivo* environments. To address this, we introduce MotorBench, an expert-annotated dataset of bacterial flagellar motors that has been curated as part of a Kaggle competition *BYU - Locating Bacterial Flagellar Motors 2025*, engaging data scientists globally to create automated detection algorithms. MotorBench and its accompanying tools are intended to serve as a benchmark for evaluating and comparing future algorithms in automated cryo-ET analysis.

## 1. Introduction

Single-celled organisms, simple as they seem, are filled with a large variety of macromolecular structures that facilitate vital cell operations such as survival, replication, and pathogenesis. Understanding these nanomachines reveals insights into how cells move, navigate, and interact with each other and their environment. One molecular machine, the flagellar motor (see Figure 1), is responsible for the motility of several pathogenic bacteria [1].

**Figure 1:**
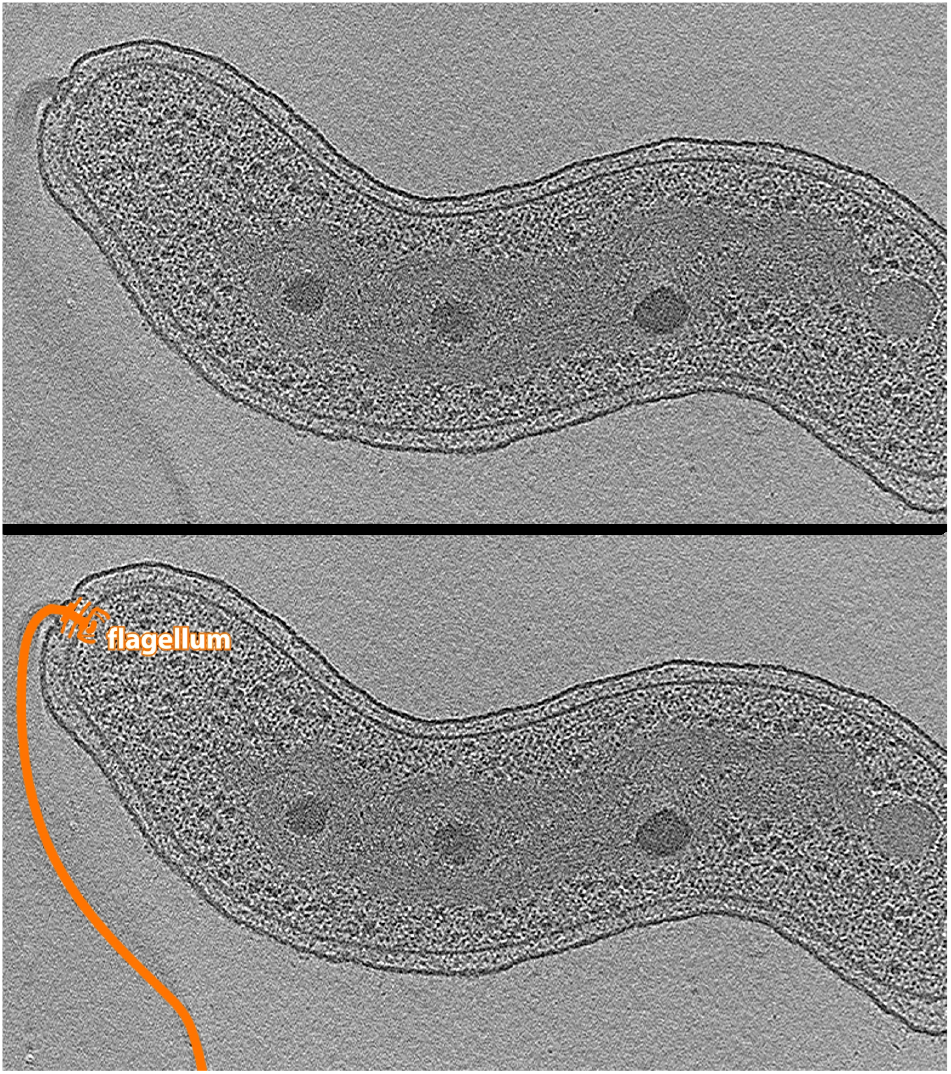
Cryo-electron tomogram of *Bdellovibrio bacteriovorus* highlighting its flagellum from *The Atlas of Bacterial and Archaeal Cell Structure* [2]. The top panel shows an unannotated slice of the cell, while the bottom panel includes an orange overlay tracing the flagellum and motor.

Flagellar motors comprise dozens of proteins spanning across the inner and outer membranes, including several ring shaped complexes that contribute to structural stability and power rotation. Among these are the stationary C-ring (in the cytoplasm) and MS-ring (in inner membrane), which drive the motion of rotor elements such as the P-ring (in the periplasm) and L-ring (in the outer membrane) [2], as seen in Figure 2. A long helical fiber, the flagellum, extends from the motor outside the cell and propels it forward. Though these macromolecular features are generally preserved across different species, the molecular makeup of flagellar motors varies considerably [3]. Because of their complexity and critical biological function, flagellar motors represent a powerful model for studying molecular machines.

**Figure 2:**
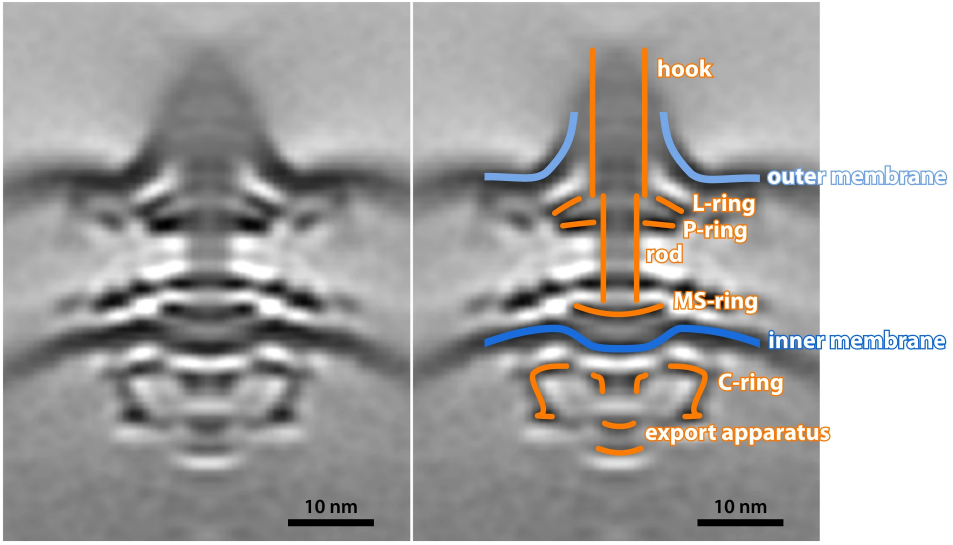
Subtomogram Averaging of a bacterial flagellar motor (left) and the same image with structural annotations (right). Key components of the motor are labeled, including the hook, rod, L-, P-, MS-, and C-rings, as well as the export apparatus. The outer and inner membranes are also indicated. This structural map highlights the complex architecture of the flagellar motor spanning multiple cellular compartments. Image from *The Atlas of Bacterial and Archaeal Cell Structure* [2].

Though bioinformatics, transcriptomics, and spatial genomics can reveal information about cellular machines, an additional way to understand how a machine works (and the way we know so much about flagellar motors) is to look at them. Structural insights into cellular machines drive progress in diagnostics, biotechnology, and precision medicine [4].

Cryogenic electron tomography (cryo-ET) is a 3D imaging technique used to visualize cellular components in a near-native state at nanometer resolution. Unlike traditional microscopy methods that require purification, crystallization, or chemical staining, cryo-ET preserves cellular architecture and context, making it particularly valuable for studying large nanomachines *in situ*.

The cryo-ET imaging workflow first requires the biological sample to be cryogenically frozen. For thin cells like bacteria, this is typically done by plunging the specimen into liquid ethane, which freezes all cellular activity so quickly that ice crystals cannot form. The sample is then inserted into the cryo-cooled vacuum chamber of an electron microscope, which uses electrons instead of light to illuminate the sample. Once inside the microscope, the sample is tilted in ∼ 3-degree increments from − 60 to 60 degrees. Each time the sample is tilted, a projection image is captured. The complete set is called a “tilt series.” The tilt series images are then combined to create a 3D reconstruction [5], or tomogram, of the specimen.

Much of the cryo-ET workflow is automated, but identifying nanomachines in tomograms is a time-intensive manual process. Tomograms have a low signal-to-noise ratio, low contrast, and limited resolution, making it difficult to identify protein structures in tomograms. In addition, radiation damage to the biological sample from incident electrons can damage the sample. Thus, samples are imaged with low doses of radiation, ultimately preserving the sample but reducing the signal in the final image. The challenges are also exacerbated by the fact that samples cannot be tilted beyond *±* 60 degrees in the microscope due to limitations of sample thickness, resulting in two wedges of information missing from the tilt series. This complicates 3D analyses [6]. Due to these limitations, locating and annotating cellular machines in tomograms requires a highly trained scientist to manually review each 3D image, a significant bottleneck in an otherwise automated workflow.

Machine learning could reduce bottleneck in cryo-ET analysis. Instead of relying on expert inspection, algorithms could be trained to detect and interpret faint structural cues embedded in tomographic data. These models excel at parsing complex visual information, even in the presence of noise and missing data, and can consistently pinpoint features of interest across large datasets. By leveraging these capabilities, researchers can streamline the interpretation of tomograms, significantly reducing the time required for annotation.

To accelerate the development of machine learning approaches, we launched a competition *BYU - Locating Bacterial Flagellar Motors 2025* on Kaggle [7], an online platform for hosting data science and machine learning competitions [8]. Using datasets provided by companies and organizations, Kaggle allows a global community of participants to come together to tackle real-world problems. Beyond competitions, Kaggle fosters collaboration by offering tools, tutorials, and a vibrant community for learning and sharing code, making it an ideal environment to accelerate scientific progress.

In addition to inspiring algorithmic innovation through competition, MotorBench and the evaluation protocol introduced here establish a standard benchmark for future methods in cryo-ET object detection, especially under *in vivo* conditions. Unlike other datasets [9, 10, 11, 12], this benchmark captures both biological complexity and imaging variability, making it suited for evaluating model robustness in real-world scenarios.

Our competition challenges participants to create a machine learning model that can detect flagellar motor centers after training on a dataset of labeled tomograms. To incentivize innovation and participation, a $65,000 prize pool will be divided between the top five teams.

Our competition launched shortly after the completion of the *CZII CryoET Object Identification Challenge* [13]. This related challenge, which was hosted on Kaggle from November 2024 to February 2025, demonstrated the effectiveness of competitions in advancing structural biology research. The challenge focused on automating the detection of five unique protein complexes in cryo-ET *in vitro* data, and it attracted over 6,800 participants from 76 countries, resulting in nearly 28,000 submissions. The winning solution [14] integrated segmentation and object detection models to identify the protein complexes, achieving a score of **0.787** (roughly interpreted as the ability to find 78.7% of the proteins).

An important difference in the datasets between the CZII’s competition and the current BYU competition lies in the cellular context of the imaged samples. The CZII tomogram dataset was composed of *in vitro* samples, that is, purified target particles were mixed with cell lysate to create a cell-like environment. In contrast, the BYU data comprises *in vivo* samples, meaning the nanomachines of interest are within living, intact cells, providing a physiologically relevant context, different from a controlled *in vitro* environments.

This fundamental difference introduces unique challenges and opportunities. *In vivo* imaging captures the complex interactions between bacteria and their host environments, necessitating advanced image analysis techniques capable of handling the variability and complexity inherent in living systems. By leveraging the proven community engagement and innovation demonstrated in the CZII challenge and numerous other competitions, we anticipate that the BYU competition will similarly attract a diverse array of participants to this new problem. Their contributions are expected to drive advancements in automated analysis of *in vivo* bacterial imaging, enhancing our ability to study bacterial behavior and interactions within their natural contexts.

## 2. Methods

### 2.1. Dataset

The tomograms collected for the competition originate from a collection curated by members of the Jensen laboratory at Caltech, who collectively acquired the raw tiltseries data for their own structural biology research [3, 15, 16, 17, 18] with no anticipation of applying machine learning techniques. Although many lab members contributed to the imaging and reconstruction of tomograms, all of the particle annotations in the training set were performed by a single researcher, Dr. Mohammed Kaplan, over an extended period of time. These annotations were primarily created to guide subtomogram averaging, which served as a form of internal validation to confirm the identity and quality of the labeled particles. Specifically, the tomograms were reconstructed using the SIRT algorithm as implemented in IMOD [19], and averaging was performed using PEET [20]. As a result, the dataset is research driven, biologically focused, and rich in contextual knowledge, but it lacks the consistency and structure typically preferred for machine learning workflows.

Because machine learning was not the original purpose, the data were stored on a shared work drive without any formal schema or relational database support. There were no accompanying metadata files or documentation, only filenames and directory names, which varied considerably in clarity and consistency. For example, early in the annotation process, Dr. Kaplan labeled several structures as “MS-rings” based on their appearance, though these were later identified as entirely different entities known as “hat structures” [15]. Tomograms were stored as .mrc files and annotations as .mod files, often co-located within directories that required manual inspection to determine correct associations. The dataset, compiled over years, brought together reconstructions from many different researchers, each with slightly different naming conventions or directory structures. While the data collection was highly collaborative, the annotation and final dataset assembly were conducted by Dr. Kaplan, whose evolving biological interpretations influenced the labels and file organization.

To make this dataset more convenient for training machine learning models, we developed a custom script to parse and standardize the directory structure, associating each annotation with its corresponding tomogram in an object-oriented format. This tool, made available through our repository tomogram datasets,[21] provides a reproducible and accessible interface to an otherwise fragmented and informally documented dataset.

The full dataset is organized into two separate directories: one with training data and one with testing data. The train directory contains subdirectories of tomogram slices stored as JPEGs, where each subdirectory contains a stack of 2D slices paired with a CSV file of training data labels.

The training data labels consist of the following elements:

- **row id**: Row index in the file.
- **tomo id**: Unique tomogram identifier (can appear multiple times if the tomogram has more than one motor).
- **Motor axis 0, 1, 2**: *z*-, *y*-, and *x*-coordinates of the motor.
- **Array shape axis 0, 1, 2**: *z, y*, and *x* dimensions of the tomogram array.
- **Voxel spacing**: The physical size of each voxel in the tomogram, given as a scaling factor in angstroms per voxel. It defines the resolution of the 3D volume by indicating how much real-world distance each voxel represents.
- **Number of motors**: Number of motors in the tomogram

The test folder stands in for the final test set, which includes a number of hidden tomograms. Due to the ongoing nature of the competition, details about the final test set cannot be disclosed until the competition ends. A submission CSV is provided to illustrate the required format, where a value of − 1 should be used for all three coordinates when no motor is present.

The raw tomogram data exhibited considerable variation in size, a result of image processing steps and the diverse hardware used to capture the images. This variability is evident across the training set, as detailed in Table 1, which lists dimensions ranging from (300, 924, 956) to (800, 1912, 1847) voxels, with a total of 817 tomograms. Such differences reflect the influence of imaging equipment specifications and subsequent computational adjustments, necessitating flexible data handling strategies to accommodate the range of sizes encountered in the dataset.

**Table 1:** Summary of Tomogram Dimensions and Counts in Training Directory.

| Tomogram Dimensions (voxels) | Train Set Count |
| --- | --- |
| (300, 960, 928) | 309 |
| (800, 928, 960) | 176 |
| (500, 924, 956) | 70 |
| (300, 959, 928) | 70 |
| (300, 928, 928) | 58 |
| (500, 928, 960) | 26 |
| (800, 928, 928) | 25 |
| (500, 1024, 1440) | 23 |
| (800, 927, 959) | 20 |
| (600, 928, 960) | 16 |
| (500, 960, 928) | 6 |
| (600, 928, 928) | 5 |
| (500, 928, 928) | 3 |
| (500, 1912, 1847) | 3 |
| (300, 924, 956) | 1 |
| (400, 960, 928) | 1 |
| (600, 960, 928) | 1 |
| (486, 1912, 1847) | 1 |
| (494, 1912, 1847) | 1 |
| (300, 956, 924) | 1 |
| (500, 927, 959) | 1 |
| **Total** | **817** |

In addition to variability in overall dimensions, voxel spacing—the physical size represented by each voxel, also varies across the training set, albeit to a lesser extent (see Table 2). Despite being subtler, this variation highlights the importance of accounting for voxel size during analysis and illustrates the challenge of developing robust machine learning models that can accommodate such differences.

**Table 2:** Summary of Tomogram Voxel Spacing and Counts in Training Directory.

| Voxel Spacing ($\text{\AA}$) | Train Set Count |
| --- | --- |
| 13.1 | 370 |
| 15.6 | 171 |
| 16.8 | 89 |
| 16.1 | 83 |
| 19.7 | 50 |
| 13.3 | 23 |
| 19.3 | 17 |
| 19.2 | 7 |
| 6.5 | 4 |
| 13.2 | 2 |
| 6.4 | 1 |
| **Total** | **817** |

Tomograms across all sets were examined and annotated using the IMOD software suite. In this manual process, the annotator visually identified each flagellar motor within the tomogram and carefully selected the precise location corresponding to the motor’s center (MS-ring region). These annotated positions were then recorded and stored as separate MOD files for further analysis. Figure 3 provides a visual example of this annotation procedure, illustrating the initial detection and final localization of the motor within the tomogram.

**Figure 3:**
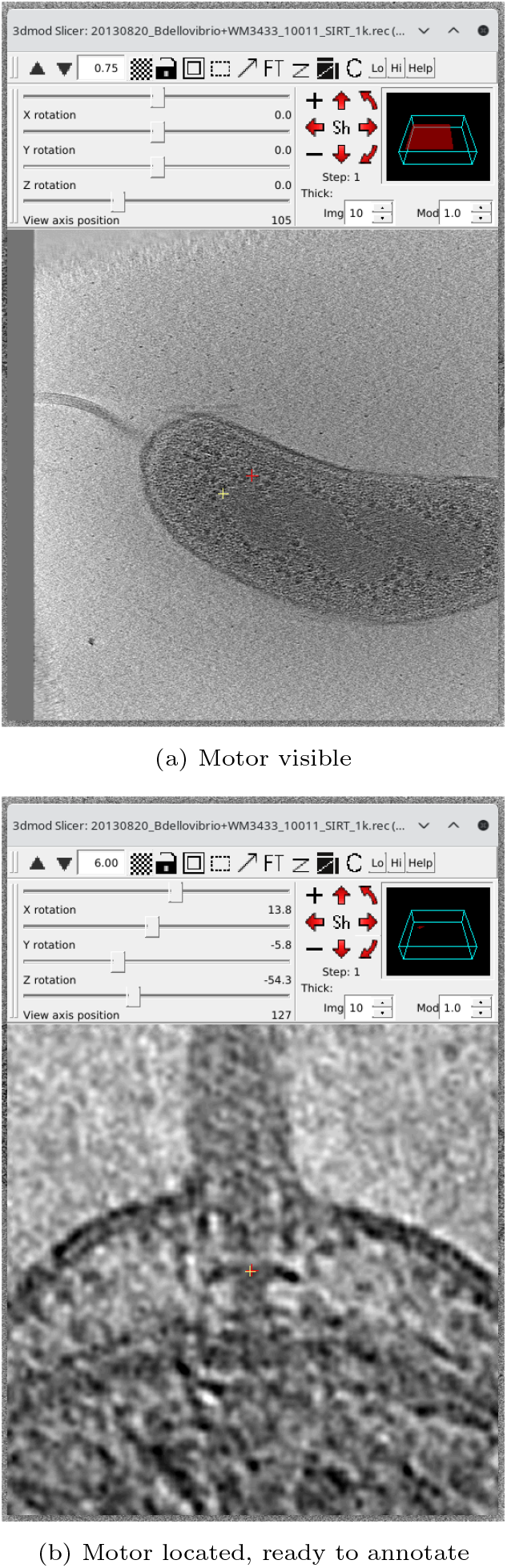
Using IMOD to find and annotate a flagellar motor in a tomogram.

Preprocessing was implemented to reduce the data size for the Kaggle competition and to normalize the images. The tomogram files (MRC) were compressed by first extracting individual slices along the *z*-axis. Each slice was clipped eliminating 1st and 99th percentiles to minimize extreme intensity values, normalized to an 8-bit (0–255) scale, and saved as a JPEG image with a 50% compression quality (see Figure 4.) The JPEG images were then organized into directories corresponding to their original tomograms. The train directory consists of 362 tomograms that contain flagellar motors (positives) and 455 tomograms without motors (negatives), providing a balanced set for training and evaluation of detection algorithms.

**Figure 4:**
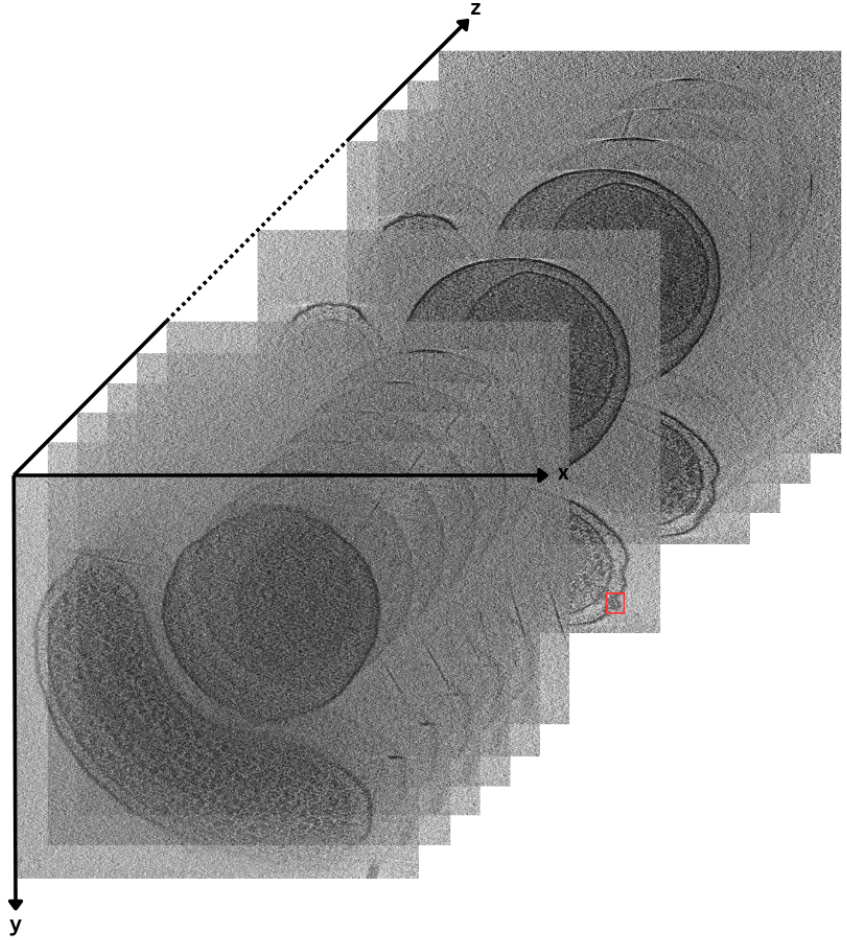
Visualization of tomogram slices from a 3D reconstruction volume, showing a series of *xy* planes stacked along the *z*-axis. Each slice reveals a cross-section of a bacterium.

Flagellar motors exhibit distinct spatial distribution patterns within the tomograms. They are notably concentrated near the center along the *z*-axis, the primary optical axis used to split tomograms into JPEG slices, as illustrated in Figure 5a. Interestingly, motors are less common near the edges along axes 1 and 2, as shown in Figure 5b. Beyond its role in the competition, the dataset’s structured format, biologically validated annotations, and opensource preprocessing tools make it a reusable benchmark for the broader machine learning and structural biology communities. By providing detailed metadata and standard evaluation formats, it supports consistent comparison across future models and approaches.

**Figure 5:**
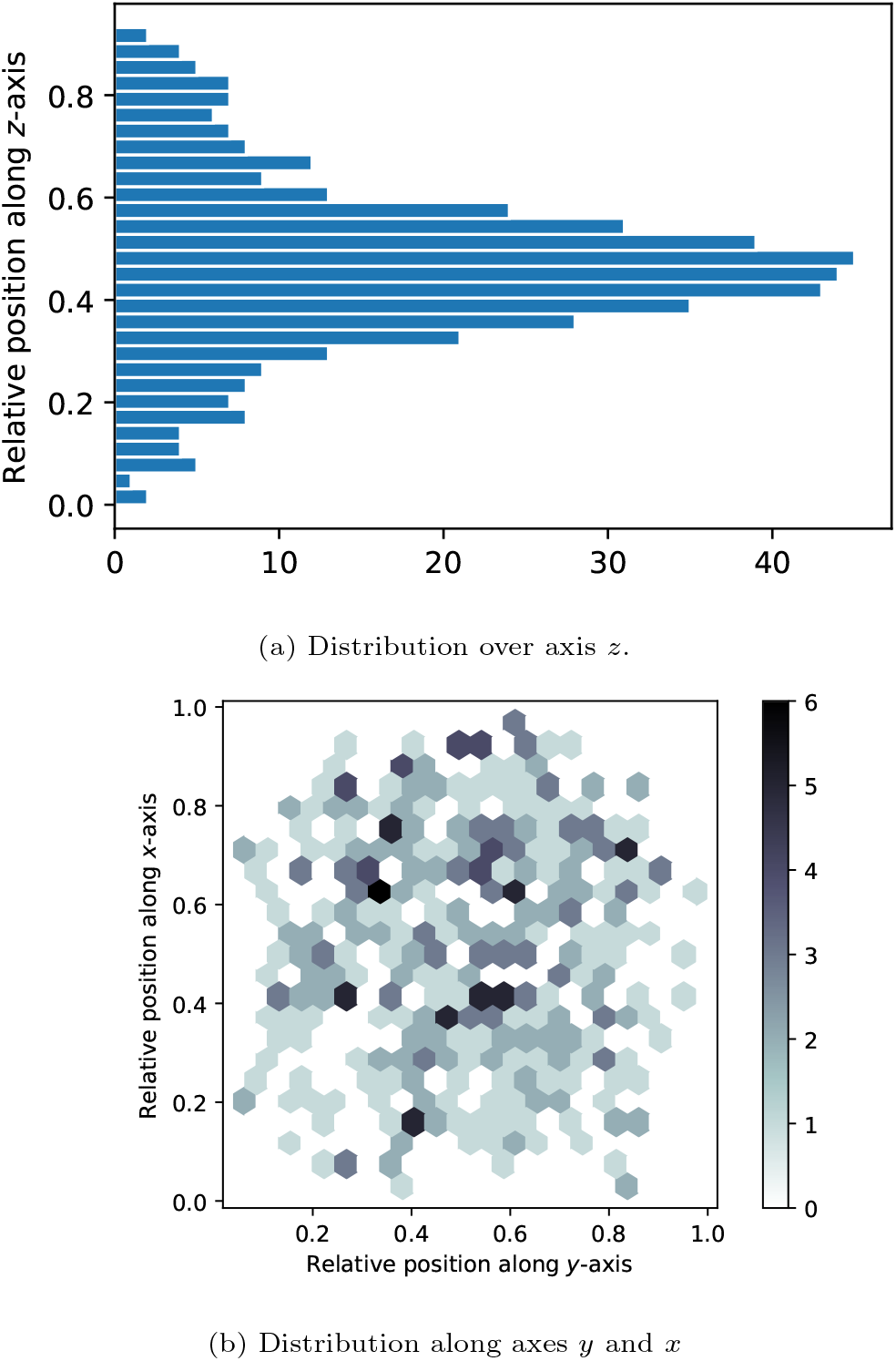
Spatial distribution of flagellar motors across different axes.

### 2.2. Evaluation Metric

Although a variety of scoring metrics were evaluated, the *F*_*β*_-score was ultimately selected for assessing the submitted models. The *F*_*β*_-score is the harmonic weighted average between precision and recall, both of which are crucial to analyzing tomograms. However, in practical applications, it would be worse for a submitted algorithm to completely miss a tomogram with a motor than it would be for a scientist to double-check false positives. Therefore, *β* = 2 is used in order to weight recall more than precision in the harmonic average.

To assess the accuracy of predicted motor locations, a preliminary evaluation step is implemented prior to computing the *F*_*β*_-score. This preliminary evaluation is applied in cases where the submitted model accurately predicts the presence of a motor. In this step, the Euclidean distance *d* = ∥*y* − *ŷ*∥ _2_ between the actual motor location *y* and the predicted motor location *ŷ* is calculated.

If *d* ≤ *τ*, where *τ* is a predefined distance threshold (set to 1000 Å), the prediction is considered correct, and the tomogram is classified as containing a motor. Otherwise, the prediction is treated as a failure to detect a motor. This misclassification subsequently influences the *F*_*β*_-score by counting the tomogram’s misclassification as a false negative.

There are four possible outcomes based on the presence or absence of a motor in both the ground truth and the model’s prediction (see Figure 6 for visual examples):

**Figure 6:**
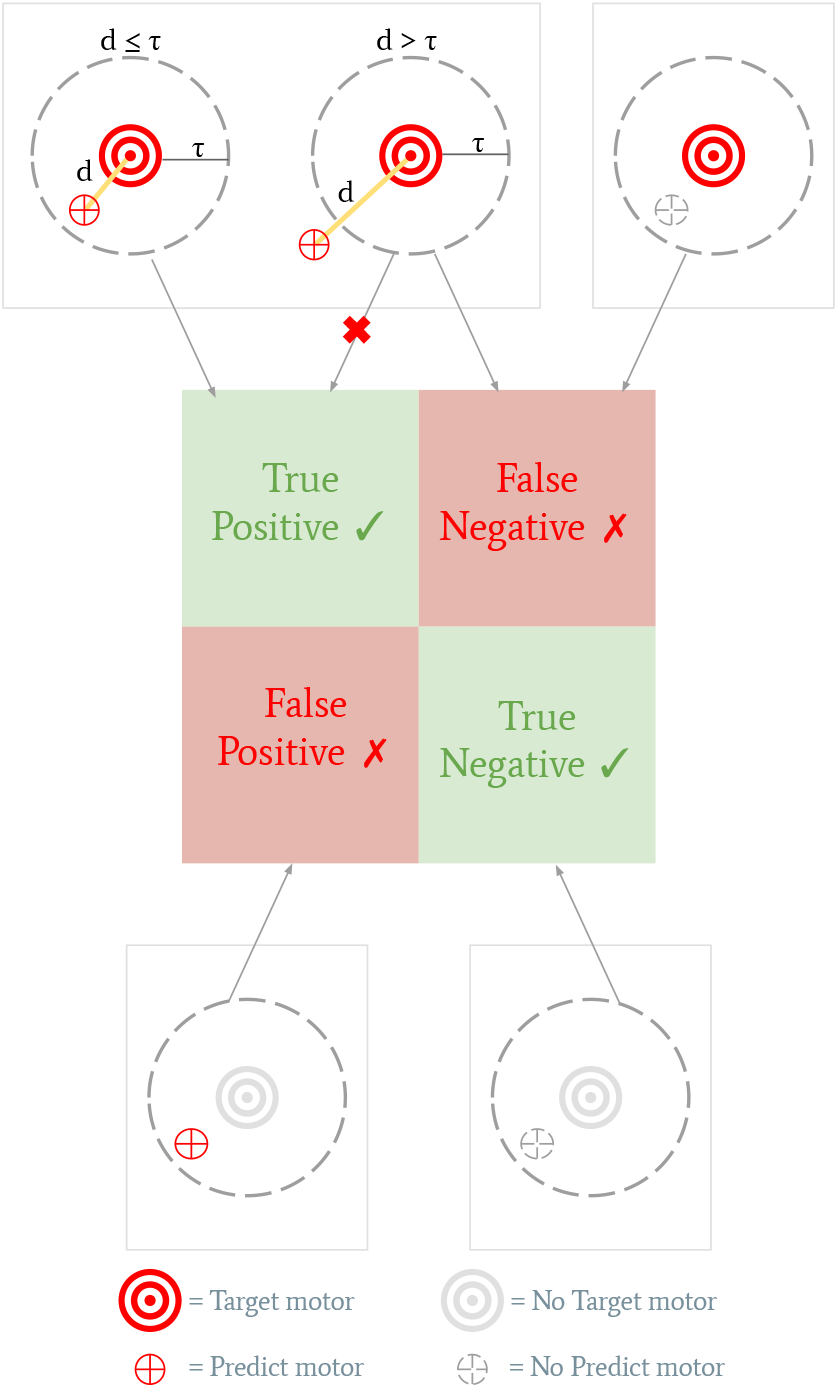
Illustration of the evaluation metric used in the competition. The primary metric is the *F*_2_-score, which emphasizes recall to reduce the risk of missing tomograms containing flagellar motors. Before computing the *F*_2_-score, a preliminary evaluation step assesses whether the predicted motor location is sufficiently close to the actual motor location. This is done by calculating the Euclidean distance *d* between the true and predicted motor positions. If *d* ≤ *τ*, where *τ* is a predefined distance threshold, the prediction is considered correct; otherwise, it is counted as a false negative. This evaluation process ensures that motor predictions are both accurate in detection and precise in localization before contributing to the final *F*_2_-score calculation. The figure visually represents this classification process.

- **True Positive (**✓): A motor is present in the tomogram, and the model correctly predicts its presence with a sufficiently accurate location (*d* ≤ *τ*).
- **False Positive (**✗): No motor is present, but the model incorrectly predicts one.
- **False Negative (**✗): A motor is present, but the model either fails to predict its presence or predicts it with an incorrect location (*d > τ*).
- **True Negative (**✓): No motor is present, and the model correctly predicts its absence.

### 2.3. In-House Notebooks and Baseline Implementation

To guide competition participants, our team created an in-house Kaggle notebook series to demonstrate possible approaches to the problem and to illustrate the submission format.

The first notebook in the series parses a subset of the provided dataset to train YOLO. For each motor annotation included in the train labels, the code extracts a subvolume of the tomogram centered around the *z*-coordinateof the motor. To form a validation set, the notebook selects a random subset of the tomograms containing 20% of the total motors. This approach avoids dataset leakage by preventing subvolumes of certain tomograms from appearing in both the training and validation set. After the split, the notebook extracts the slices from each subvolume and annotates them with a bounding box corresponding to the *x*- and *y*-coordinates of the train labels.

The second notebook allows participants to visualize the preprocessing techniques employed in the first notebook by displaying a small selection of images and their corresponding motor locations (see Figure 7).

**Figure 7:**
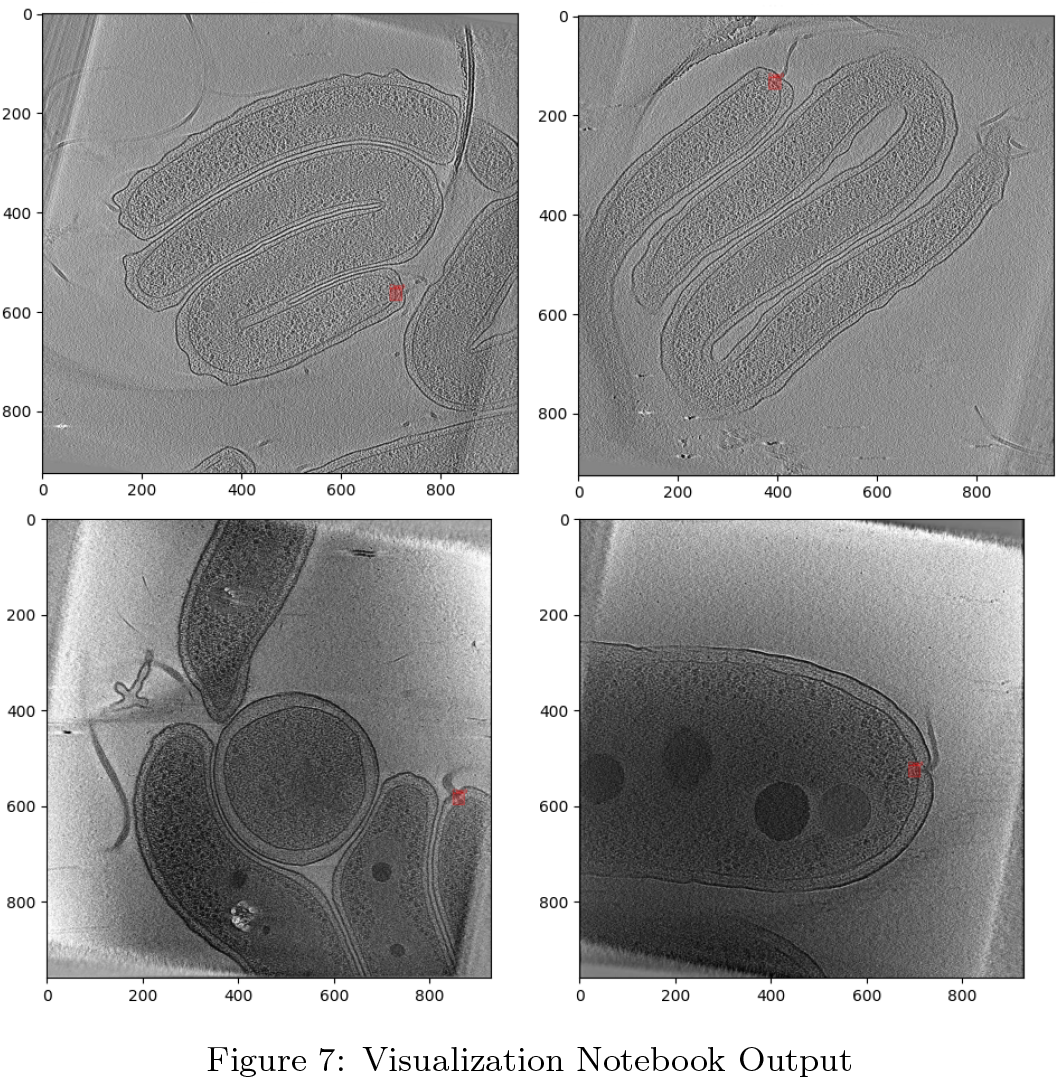
Visualization Notebook Output

With the dataset properly formatted using the parsing notebook, a third notebook fine-tunes a pretrained YOLOv8 model to detect flagellar motors. Training proceeds for up to 30 epochs, with early stopping triggered if the validation loss does not improve over five consecutive epochs. The optimization is guided by the distance-focal loss (DFL) function [22], and model checkpoints are saved every five epochs. The model achieving the lowest validation DFL loss is ultimately selected as the final version.

To generate a submission in the proper format, the final notebook loads the model weights and the Ultralytics library in an offline setting to reproduce the same evaluation pipeline as before, but in a competition-specific format. Upon submission to the competition, the notebook was granted access to the true public test set and executed the evaluation on each provided tomogram, generating a CSV file containing the predictions. When processed by Kaggle, the generated predictions achieved an *F*_*β*_-score of 0.457 according to the evaluation metric.

## 3. Conclusion

Understanding the mechanical function of bacterial nano-machines remains a key challenge in both fundamental biology and the pursuit of novel therapeutics. Cryo-ET provides a view of these intricate structures within their native cellular contexts; however, locating and annotating machines inside tomograms continues to be a major hurdle in the imaging workflow.

The launch of a Kaggle competition focused on *in vivo* tomograms offers a new avenue for advancing automated nanomachine detection. This dataset not only reflects the biological complexity of living systems but also presents machine learning researchers with a problem that is both timely and technically demanding.

By encouraging collaboration between biologists and data scientists worldwide, the competition may lead to faster models capable of managing noisy, variable, and real-world data. As submitted solutions continue to evolve, improvements in accuracy, robustness, and practical applicability are expected alongside tools that may help accelerate discovery across microbiology, structural biology, and biomedical imaging. Scalable, intelligent solutions are essential to the future of high-throughput cryo-ET analysis, and this competition represents an early but meaningful step in that direction.

Upon the conclusion of the competition, both the training and test datasets will be publicly available on Kaggle [7] and the CZI CryoET Data Portal [23]. As the dataset, metric, and tools remain openly accessible beyond the competition timeline, they provide a reproducible benchmark for evaluating new approaches in cryo-ET image analysis. We hope this benchmark will catalyze longterm progress and become a standard reference point for future algorithmic work in the field.

## 3.1. Acknowledgments

Cryo-ET imaging in the Jensen lab was supported by NIH Grant RO1 AI127401 and other sources. We acknowledge the Brigham Young University Office of Research Computing (ORC) for providing the computational resources that supported this work. We also thank the College of Computational, Mathematical, and Physical Sciences at BYU for supporting undergraduate student participation in this research. We are especially grateful to Kaggle for contributing $50,000 to the competition prize pool and for their collaborative support throughout the project. In particular, we thank Walter Reade, Maggie Demkin, and Elizabeth Park at Kaggle for their invaluable assistance and guidance. This research was funded in part by the Chan Zuckerberg Initiative (CZI). Mohammed Kaplan acknowledges funding from the University of Chicago (startup package).

## Notes

### Competing Interest Statement

The authors have declared no competing interest.

